# Mechanism of riboregulation of p62 protein oligomerisation by vault RNA1-1 in selective autophagy

**DOI:** 10.1101/2021.04.12.439413

**Authors:** Magdalena Büscher, Rastislav Horos, Kevin Haubrich, Nikolay Dobrev, Florence Baudin, Janosch Hennig, Matthias W. Hentze

## Abstract

Macroautophagy ensures the clearance of intracellular substrates ranging from single ubiquitinated proteins to large proteotoxic aggregates and defective organelles. The selective autophagy receptor p62 binds these targets and recruits them to double-membrane vesicles, which fuse with lysosomes to degrade their content. We recently uncovered that p62 function is riboregulated by the small non-coding vault RNA1-1. Here, we present detailed insight into the underlying mechanism. We show that the PB1 domain and adjacent linker region of p62 (aa 1-122) are necessary and sufficient for specific vault RNA1-1 binding, and identify lysine 7 and arginine 21 as key hinges for p62 riboregulation. Chemical structure probing of vault RNA1-1 further reveals a central flexible loop within the RNA that mediates the specific p62 interaction. Our data define molecular determinants that govern mammalian autophagy via the p62-vault RNA1-1 riboregulatory pair.

## INTRODUCTION

The human autophagy receptor p62 (also known as SQSTM1, ZIP or ORCA) guides selective macroautophagy of intracellular cargo to maintain homeostasis in situations of proteotoxic stress or starvation [1–3]. Following activation, p62 oligomerizes and forms large protein assemblies – so-called sequestosomes – that are linked to the elongating autophagic membrane [4]. p62 recognises ubiquitinated cargo via its C-terminal ubiquitin-associated (UBA) domain [5] and N-arginylated proteins via its ZZ-type zinc finger domain, respectively [6]. The central LC3-interacting region (LIR) binds the ATG8-like proteins LC3 and GABARAP, thereby bridging intracellular cargo to the autophagic membrane [7].

Since the affinity of p62 monomers for specific cargo and the ATG8-like proteins is rather modest, protein oligomerisation is essential to achieve high avidity protein interactions while maintaining selectivity [8]. Oligomerisation is mediated by the N-terminal Phox and Bem1 (PB1) type I/II domain that comprises an OPCA motif (short for OPR: octicosapeptide repeat, PC: Phox and Cdc motif, AID: Atypical protein kinase C interaction domain) and a conserved lysine which can align in a head to tail-fashion [9–11]. Strikingly, mutations that disrupt PB1 domain-mediated oligomerisation prevent p62 engagement in autophagy and thereby highlight the ‘effector’ function of PB1-mediated oligomerisation in this process [1,4,11–13]. Intrinsic modulators of p62 oligomerisation include the formation of stabilising disulphide bonds between cysteine 105 and 113 upon oxidative stress or ZZ domain binding [6,14], as well as inhibitory posttranslational modifications within the oligomerisation interface [15,16]. Besides, the linker region between the PB1- and ZZ domain of p62 (aa 100-113) was shown to auto-regulate p62 by interaction with the ZZ domain [17].

We recently discovered that the small non-coding vault RNA1-1 directly binds to p62 and regulates p62 oligomerisation and hence function in autophagy [18,19]. Vault RNAs are transcribed by RNA polymerase III and were originally identified more than three decades ago as components of the so-called vault particle [20]. The four human vault RNA paralogs share almost identical sequences at their 3’ and 5’ ends while their central domains vary in sequence and length [21,22]. We found that vault RNA1-1 is the prime p62-interacting vault RNA, but the molecular determinants of their specific interaction and the mechanism of how these control oligomerisation remained unresolved [19]. We previously described a ZZ domain mutant (R139A/K141A) and a PB1 domain mutant (R21A/D69A/D73A) that both exhibit reduced RNA binding while showing increased or diminished oligomerization, respectively [19]. These findings implicated both domains in riboregulation, suggesting that the PB1 domain itself or PB1 domain-mediated oligomerization may be required for p62’s RNA binding activity [19]. Here, we delineate the key determinants that mediate binding and specificity of the p62 -vault RNA1-1 interaction, uncovering dependencies between oligomerization and RNA binding. Our results elucidate the molecular details of this prime example of riboregulation.

## MATERIAL AND METHODS

### Antibodies and reagents

Antibodies and reagents used in this study are listed in Supplementary Table 1.

### Cell lines and culture conditions

HuH-7 cells (derived from a hepatocellular carcinoma of a human aged 57 years) and their derivatives were cultured in DMEM containing 1 g/l glucose supplemented with 10% heat-inactivated FCS (Fetal Calf Serum, Gibco, Cat#: 10270-106), 2 mM L-glutamine (Thermo Scientific, Cat#: 25030081) and 100 U/ml Penicillin/Streptavidin (Thermo Scientific, Cat#: 15140122) at 37°C and 5% CO_2_. The cells were routinely passaged 2-3 times per week. For this purpose, the cells were dissociated through the addition of Trypsin-EDTA (0.05%, Gibco, Cat#: 25300-054). HuH-7 Flp-In cell line stocks and their derivatives were cultured in medium containing zeocin (100 µg/ml, InvivoGen, Cat#: ant-zn-05) or hygromycin B Gold (200 µg/ml, InvivoGen, Cat#: ant-hg-1) depending on expressed marker genes. The parental hepatocellular carcinoma HuH-7 Flp-IN cell line (one FRT integration site; Clone C2111) was first established in [23], the HuH-7 p62 KO cell line in [19].

### Cloning

Restriction-free cloning was performed according to [24].

### CRISPR/Cas9 genome editing of cell lines

CRISPR/Cas9 genome editing was based on [25,26]. In brief, guide RNAs targeting the vtRNA 1-1 locus (Supplementary Table 2, [19]) were predicted using the CRISPOR online tool (http://crispor.tefor.net; Version May 2017), the highest-scoring candidates ordered from Sigma-Aldrich, annealed and ligated into BbsI linearised pSpCas9(BB)-2A-GFP/RFP/Cer constructs (Kindly provided by the Noh Laboratory, EMBL Heidelberg). Combinations of the generated plasmids were nucleofected into HuH-7 Flp-IN cells using the SF Cell Line 4D-Nucleofector X Kit according to the manufacturer’s guidelines, with 1Mio cells and 1µg of total plasmid DNA in a 100 µl setup running programme FF137. As a negative control, a mixture of all parental plasmids was used. Fluorescence-activated single-cell sort of double/triple-positive cells was performed 48h after nucleofection. Upon expansion, the clones were tested for vault RNA 1-1 deletion by PCR from genomic DNA using locus spanning primers (Supplementary Table 2). Cell lines lost the transiently transfected plasmids after outgrowth as checked via FACS.

### Transfections and Treatments

Transfections were performed using Lipofectamine 3000 according to the manufacturer’s guidelines. For cell-based assays, XIE62-1004-A (50mM in DMSO; [19]), was diluted in PBS to a concentration of 2.5 mM and added to the medium in a final concentration of 10 µM.

### Native immunoprecipitation (IP)

For IPs one confluent Ø 15cm dish per IP served as starting material. The cells were washed twice with ice-cold PBS. The buffer was aspirated completely and the cells lysed on ice through the addition of 750 µl Trit-Lysis buffer (20 mM Tris-HCl pH7.4, 150 mM NaCl, 1 mM EDTA, 1 mM EGTA, 1 % Triton X-100) supplemented with cOmplete protease inhibitor tablets (Roche, Cat#: 11873580001) and 5 µg/ml RNAsin (Promega, Cat#: N2511). The cells were collected through scraping and homogenised by pipetting. Lysates were cleared by centrifugation for 10 min at 13 000xg, 4°C. The supernatant was transferred into a DNA LoBind 1.5 ml reaction tube (Eppendorf), the protein concentration determined by Bradford assay and the input material adjusted accordingly (1-2 mg/IP). Per IP, 25µl anti-HA magnetic bead slurry was washed twice with PBS prior to addition to the samples. The IPs were performed for 1-2 h at 4°C with constant rotation. Following, the samples were washed six times with 1ml of Trit-Lysis buffer and the reaction tubes exchanged after every second wash. For pH elution 50µl of 0.1M glycine pH2 were added per condition and incubated for 5min at room temperature. The eluates were transferred into a new reaction tube and neutralised through addition of 7.5 µl of 1 M Tris-HCl pH8.5.

For protein analysis, 1% of input material and 15-20% of elution were analysed by Western blotting. For RNA analysis 5% of input material and 70% of elution were processed by the addition of 400 µl RNA lysis buffer (Zymo) and RNA extraction using the Zymo Quick-RNA™ MicroPrep RNA extraction kit (Zymo) following the manufacturer’s guidelines including the DNaseI on-column DNA digest. Elution was performed with 15 µl RNase-free H_2_O.

### Polynucleotide kinase (PNK) assay of FLAG-HA tagged proteins

For the PNKs one 80-90% confluent Ø10cm or Ø15cm dish per condition served as starting material. The cells were washed twice with ice-cold PBS, the PBS was aspirated completely and the cells were UV-crosslinked at 150 mJ/cm^2^. Subsequently, the cells were lysed on ice with 0.75-1ml PNK lysis buffer (50 mM Tris-HCl pH7.4, 100 mM NaCl, 0.1 % SDS, 1 mM MgCl_2_, 0.1 mM CaCl_2_, 1 % NP40, 0.5 % sodium deoxycholate supplemented with cOmplete protease inhibitors) and transferred into 1.5ml reaction tubes. The lysates were homogenized by sonication on ice (Branson cell disruptor B15: 3x 10 s, 50% amplitude, level 4) and cleared by centrifuged at 16 000xg for 10 min, 4°C. The protein concentration was measured by Bradford assay and the input material adjusted accordingly (1-2 mg/condition). Following the lysates were treated with 5 ng/µl RNase A and 2 U/ml Turbo DNase for 15 min at 37°C and 1100 rpm. Per condition, 25µl anti-HA magnetic bead slurry was washed twice with PBS prior to addition to the samples. The IPs incubated for 1.5-2 h at 4°C under constant rotation. Subsequently, the IPs were washed three times with PNK lysis buffer and three times with PNK wash buffer (50 mM Tris-HCl pH7.4, 50 mM NaCl, 10 mM MgCl_2_, 0.5 % NP-40, supplemented with cOmplete protease inhibitors). Radioactive labelling of retained RNA was performed on beads in PNK wash buffer containing 0.1 µCi/µl [γ-^32^P] ATP, 1 U/µl T4 PNK and 1 mM DTT for 15 min at 37°C and 850 rpm. After another four washes with PNK wash buffer, the proteins were eluted by addition of 50 µl 0.1 M glycin pH2 for 5 min and eluates neutralized with 7.5 µl 1M Tris-HCl pH8.5. The samples were complemented with 4x sample buffer containing 200 mM DTT, heated to 70°C for 3 min, resolved by SDS-PAGE and blotted onto nitrocellulose membranes. The membranes were rinsed, dried and radioactive signal detected with a phosphorimaging screen for 3-72 h. Following, the membrane was used for Western blot analysis. Image analysis was performed with Fiji Image J 2.0.0 ([31], https://imagej.net/Fiji). The relative PNK signal was calculated and normalised to the relative Western blot signal to account for variation in IP efficiency.

### SDS-PAGE and Western blotting

For standard protein analysis cells were washed twice with ice-cold PBS, the buffer aspirated completely and cells lysed through the addition of RIPA lysis buffer (25 mM Tris-HCl pH7.6, 150 mM NaCl, 1% NP-40, 1% sodium deoxycholate, 0.1% SDS supplemented with cOmplete protease inhibitor and 0.01 U Benzonase). Following the cells were collected by scraping and protein concentration measured by Bradford assay. The lysates were mixed with 4x NuPage LDS Sample buffer supplemented with 200 mM DTT before denaturation at 70°C for 3min. Typically 5-15 µg total protein were resolved by SDS-PAGE on 4-15% TGX Precast gels using 1x Laemmli running buffer (25 mM Tris, 192 mM glycine, 0.1%SDS pH 8.3). The proteins were transferred onto PVDF or nitrocellulose membranes using the Trans-Blot Turbo Transfer System (Bio-Rad) and transfer efficiency assessed by Ponceau Red staining. The membranes were blocked in PBS-T 5% milk (1xPBS, 0.1% Tween 20, 5% w/v milk powder) for 1 h at room temperature. Primary antibodies were diluted in PBS-T 5% milk and added to the membrane overnight at 4°C under constant shaking. After three PBS-T (1xPBS, 0.1% Tween 20) washes, each for 5 min, the membrane was incubated with HRP-conjugated secondary antibody in PBS-T 5% milk for 1 h at room temperature. Following three PBS-T washes the western blots were developed using ECL on a BioRad ChemiDoc MP Imaging system with auto-capture function.

### Northern blot

Typically, 15-20 µg of total RNA were mixed with 2x RNA gel loading dye (95% formamide; 0.025% xylene cyanol and bromophenol blue; 18 mM EDTA; 0.025% SDS), denatured for 5 min at 95°C, loaded onto a denaturing 8 % polyacrylamide gel (8% Acrylamide/Bis 19:1, 6 M Urea, 0.5x TBE) and separated for 1-2 h at 350 V in 0.5xTBE. A semi-dry blotting apparatus was used to transfer the RNA onto a Hybond N^+^ membrane with 0.8 mA/cm^2^ for 2.5 h in 0.5xTBE. Following RNA and membrane were crosslinked by UV exposure at 150 mJ/cm^2^.

The membrane was pre-hybridised (5xSSC, 7% SDS, 20 mM NaPi, 1x Denhardt solution, 0.1 mg/ml salmon sperm DNA) for 1 h at 50°C before the addition of a radioactively ^32^P-labelled DNA antisense oligonucleotide probe for overnight incubation at 50°C (Supplementary Table 2). The DNA probe was labelled with PNK and purified over a Chroma Spin chromatography column according to the manufacturer’s guidelines. Finally, the membrane was washed three times with a high stringency buffer (5x SSC, 2% SDS) and three times with a low stringency buffer (1x SSC, 1% SDS) before visualising the radioactive signal by 4-16 h exposure to a phosphorimaging screen.

### Protein expression and purification of recombinant full-length MBP-p62 and mutants thereof

Protein expression was performed according to [27].

### Protein expression and purification of recombinant p62 (1-122, D69A/D71A/D73) for structural studies

The oligomerisation deficient p62 truncation p62(1-122, D69A/D71A/D73A) was cloned into pCoofy4 (Ref: PMID: 23410102) using restriction-free cloning [24], transformed into the Rosetta 2(DE3) strain and expressed in M9 medium with ^15^NH_4_Cl as the sole nitrogen source at 22°C O/N upon induction with 0.5 mM IPTG. The cells were lysed in a buffer containing 50 mM Tris pH 8, 750 mM NaCl, 10% glycerol, 20 mM imidazole and cOmplete, EDTA-free Protease Inhibitor Cocktail using a microfluidizer (M-110L, Microfluidics Inc., USA). Subsequently, the protein was purified from the cleared lysate by Nickel affinity chromatography (HisTrap, Cytiva) in the same buffer. The affinity tag was removed by 3C protease cleavage (EMBL, PepCore), dialysis and an additional passage over the HisTrap column. As a final purification step, oligomers were removed by gel filtration on a Superdex S75 10/300 in a buffer containing 20 mM MES pH 6.5, 100 mM NaCl, 0.2 mM TCEP and 0.05% NaN_3_.

### Nuclear Magnetic Resonance Spectroscopy

NMR spectra were acquired on a Bruker Avance III spectrometer with a cryogenic triple-resonance probe and field strength of 18.8 T corresponding to proton Larmor frequency of 800 MHz) at 298 K. For NMR titrations protein samples at 80 µM concentration in 20 mM MES, pH 6.5, 100 mM NaCl, 0.2 mM TCEP, 0.05 % NaN_3_ were titrated with full-length vault RNA 1-1 in the same buffer. At each titration step, an apodization-weighted sampled HSQC was collected [28]. Spectra were processed using NMRPipe [29] and analysed using NMRFAM SPARKY [30]. The yields and solubility of this construct did not enable sufficient signal-to-noise ratio for backbone assignments and subsequent mapping of chemical shift perturbations onto the structure.

### In vitro transcription of RNA

For biochemical assays: pUC57-T7-vaultRNA1-1 plasmids were linearized and used for in vitro transcription of RNA using the MEGAshortscript kit (AM1354, Thermo Fisher) with ^32^P-αUTP (SRP-210, Hartmann) according to the manufacturer’s guidelines. RNA was gel purified and phenol-chloroform extracted, dissolved in water and its concentration measured by QuBit assay (Thermo).

For NMR: pUC57-T7-vaultRNA1-1 plasmids were linearized and used for in vitro transcription of RNA in a large-scale reaction containing 100 mM HEPES-KOH pH 7.5, 10 mM MgCl_2_, 2 mM Spermidine-HCl, 40 mM DTT, 0.1mg/ml BSA, 7.5 mM each NTP, 800 units/ml RNasin, 10 units/ml IPP, 25 µg/ml template DNA and 10 000 units/ml T7 RNA polymerase and incubated 8h at 37°C. The RNA was gel purified, extracted by electrophoresis, precipitated and its concentration measured by Nanodrop.

### Electromobility shift assay (EMSA)

Before the reaction, RNA was denatured 2 min at 95°C and subsequently refolded in the presence of 2.5 mM MgCl_2_. The EMSA reactions contained typically 10 nM-2 mM protein, 10 nM radioactively labelled RNA (150 fmol, 3 kcpm), 150nM non-labelled bacterial tRNA, 1 mg/ml of BSA, 10 mg/ml RNAsin, 5 mM DTT, 0.5 mM PMSF, 2.5 mM MgCl2, 100 mM KCl; 20 mM HEPES pH7.9; 0.2 mM EDTA and 20% glycerol. In case of competitive EMSAs the binding reaction was competed with non-labelled RNA at a typical range of 0.1 to 2µM that was added to the protein together with the labelled probe. Reactions were incubated 10 min at room temperature, loaded onto a native 5% acrylamide gel and ran overnight at 70 V in 0.5xTBE. Subsequently, the gel was dried for 1 hour at 80°C and signal visualized by exposure to a phosphorimaging screen.

### RNase footprinting

*In vitro* transcribed RNA was dephosphorylated with Fast alkaline phosphatase (FastAP) according to the manufacturer’s guidelines and purified over a Zymo Quick-RNA Miniprep column. Subsequently, the RNA was radioactively labelled in a T4 PNK reaction with ^32^P-γUTP (Hartmann) and purified over a Chroma Spin chromatography column. Prior to RNase footprinting, the RNA was denatured for 10 min at 65°C and allowed to refold by ramping down 1°C/30s to room temperature. Per condition 8µl containing 20 µM 3’-end ^32^P-labelled RNA (∼1600 cps) and 6 µM protein were assembled in Binding buffer (20 mM HEPES pH8, 100 mM KCl, 0.01% NP-40, 5% Glycerol, 2.5 mM MgCl_2_, 1 mM DTT supplemented with cOmplete proteinase inhibitor (EDTA-free)) and incubated for 10 min at room temperature. RNase A (5 ng/µl) was prediluted (1:200 000, 1:500 000) in H_2_O and 2 µl dilution added per reaction. The digest was incubated for 15 min at room temperature. Following the RNA was extracted by Phenol-Chloroform extraction using TRI Reagent and dissolved in 2x RNA Gel loading dye. For the alkaline ladder, 2 µl containing 20 µM 3’-end ^32^P-labelled RNA (∼400 cps) were mixed with 8 µl 50 mM NaHCO_3_ pH 9.2, incubated at 95°C for 5 min and supplemented with RNA gel loading dye. The samples and ladder were loaded onto a denaturing 8% polyacrylamide gel (8% Acrylamide/Bis 19:1, 6M Urea, 0.5x TBE) and separated at 42 W (50-55°C) in 0.5xTBE. The gel was fixed for 10 min in a bath of 6% acetic acid and 10% EtOH, dried on Whatman filter paper in a vacuum gel drier at 80°C for 40 min and the radioactive signal visualised by exposure to a phosphorimaging screen.

### Chemical structure probing of RNA

Per compound four probing reactions were performed with one untreated control and three treated samples of varying concentrations. Therefore, 20 pmol *in vitro* transcribed RNA were diluted in 12 µl H_2_O, denatured at 95°C for 2 min and directly transferred on ice. The RNA was allowed to fold for 20 min at 37°C upon addition of 12 µl 2x Folding buffer (40 mM HEPES pH8, 300 mM KCl, 0.02% NP-40, 10% glycerol, 5 mM MgCl_2_). Subsequently, 12 µl protein [15 µM] in 1x Folding buffer was added to the reaction and incubated for 10 min at 37°C before splitting the reaction into 4x 8 µl each. For the modification reactions, 92 µl Mod. buffer A for DMS (20 mM HEPES pH7, 150 mM KCl, 0.01% NP-40, 5% Glycerol, 2.5 mM MgCl_2_) or Mod. buffer B for CMCT (20 mM HEPES pH8, 150 mM KCl, 0.01% NP-40, 5% Glycerol, 2.5 mM MgCl_2_) were added. For DMS treatments 1µl of 16x, 4x or 2x DMS diluted in EtOH was added to the reactions, immediately mixed and incubated 15 min at 37°C. For CMCT treatments 5µl, 10µl or 15µl of CMCT solution (42 mg/ml in H_2_O) were added to the reactions, immediately mixed and incubated 15 min at 37°C. The reactions were stopped on ice, RNA extracted with TRI reagent according to the manufacturer’s guidelines and EtOH precipitated. Modified residues were detected by primer extension reaction using a radiolabelled DNA oligonucleotide (T4 PNK reaction and purification over a Chroma Spin 10 chromatography column) complementary to the 3’-end of the RNA. For primer extension, 1µl of 10µM radiolabelled primer was annealed to 2µl of extracted RNA and used in a 5µl reaction with AMV reverse transcriptase according to the manufacturer’s guidelines. For the sequencing lanes *in vitro* transcribed RNA was used in primer extension reactions that contained respective ddNTPs. The reactions were stopped with 20µl Stop buffer (50 mM Tris pH8.3, 0.5% SDS, 7.5 mM EDTA) and the cDNA EtOH-precipitated and separated on a denaturing 8% polyacrylamide gel (8% Acrylamide/Bis 19:1, 8M Urea, 0.5x TBE) at 42W in 0.5xTBE. Subsequently, the gel was fixed for 10 min in a bath of 6% Acetic acid and 10% EtOH, dried on Whatman filter paper in a vacuum gel drier at 80°C for 40 min and the radioactive signal visualised through exposure to a phosphorimaging screen.

### Reverse transcription and quantitative PCR (RT-qPCR)

Reverse transcription was performed with 500 ng of total RNA or 7 µl of purified RNA from IPs in a 10 µl reaction using the Maxima First-strand synthesis kit according to the manufacturer’s guidelines. Control reactions that do not include the reverse transcriptase (-RT) were included for each sample. The obtained cDNA was diluted 1:20-1:50 in nuclease-free water and assembled in 10 µl qPCR reactions with 5 µl cDNA, 4.6 µl Fast SB SYBR Green Master Mix, 0.2 µl each primer (10 µM). Primer sequences can be found in Supplementary Table 2. The analysis was performed on an Applied Biosystems QuantStudio 6 Flex Real-Time PCR system in the default standard two-step cycling run. Relative expressions were calculated with the ∆∆CT method. The mean CT value was used when normalizing to multiple housekeeping genes.

## QUANTIFICATION AND STATISTICAL ANALYSIS

Statistical analysis was performed with GraphPad Prism 8.3.1 (279; www.graphpad.com). Details for each experiment can be found in the corresponding figure legend. Image analysis was performed with Fiji Image J 2.0.0 ([31], https://imagej.net/Fiji). Structure representations were generated using PyMOL Molecular Graphics System 2.0.7 (https://pymol.org). NMR analysis was performed with NMRPipe ([29], https://www.ibbr.umd.edu/nmrpipe/) and NMRFAM SPARKY ([46], https://nmrfam.wisc.edu/nmrfam-sparky-distribution/).

## RESULTS

### The PB1 domain and adjacent linker region of p62 mediate specific vault RNA 1-1 binding

To identify the protein region that is required for RNA binding in a systematic manner, we generated a series of FLAG-HA tagged p62 truncation constructs (Figure 1A), and assessed their RNA-binding capacity by polynucleotide kinase (PNK) assays in p62 knockout (KO) HuH-7 cells (Figure 1B). UV-crosslinking introduces covalent bonds between p62 and bound RNAs, the cells are lysed, and lysates subjected to limited RNase treatment before p62 immunoprecipitation and end-labelling of crosslinked RNA with PNK (Figure 1B). The eluates are resolved by SDS-PAGE and RNA binding is assessed by phosphorimaging and Western blotting, respectively. Although UV-crosslinking is not considered to promote protein-protein crosslinking in general [32– 34], we noticed earlier that p62 oligomers can be UV-crosslinked, allowing the assessment of oligomerization in addition to RNA binding[19].

**Figure 1:**
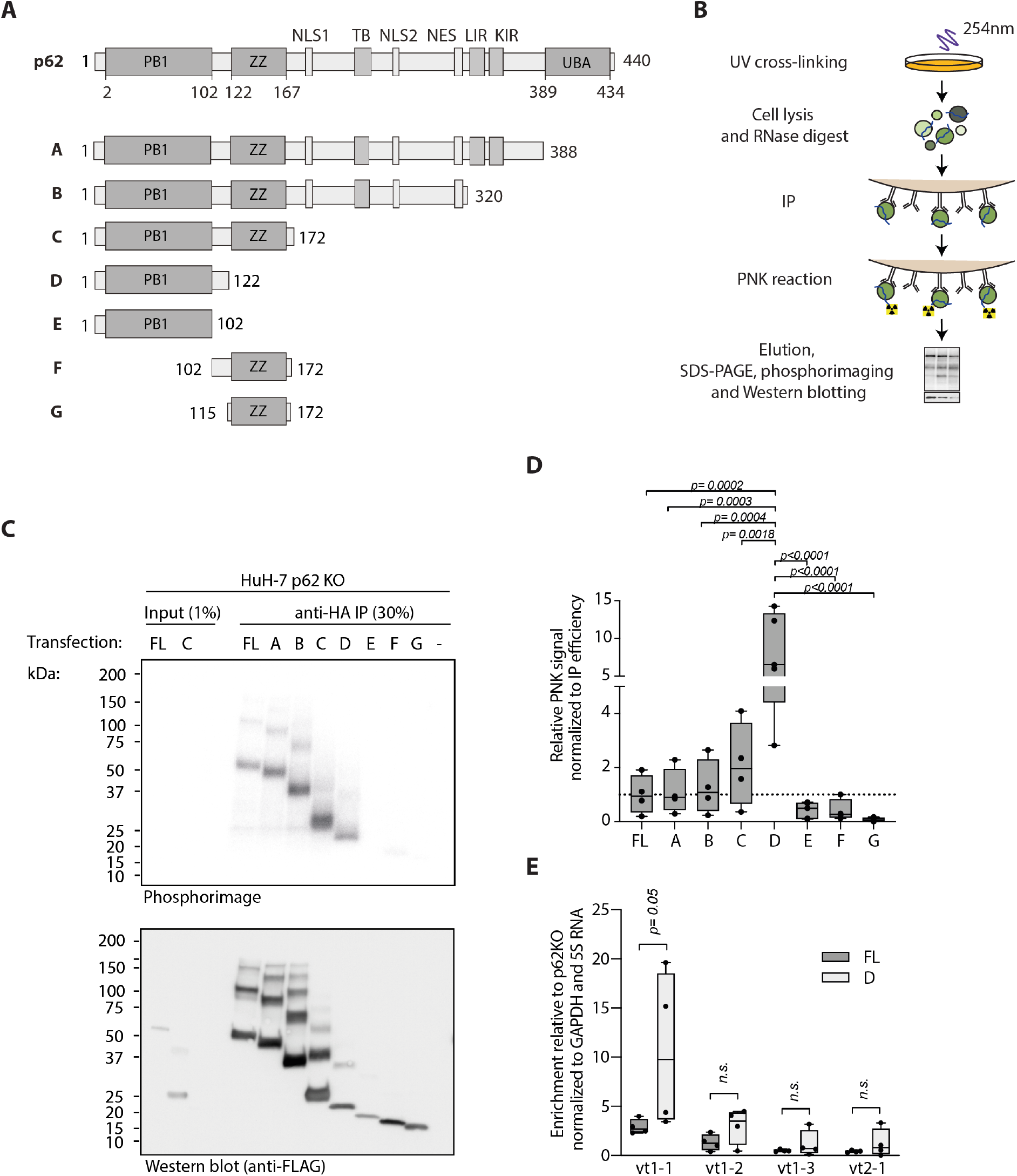
The PB1 domain and adjacent linker region are necessary and sufficient for p62’s RNA binding capacity and specificity towards vault RNA1-1. (A) Schematic overview of full-length p62 and truncation constructs with N-terminal FLAG-HA tags. (FL: full-length p62; PB1: Phox and Bem1; ZZ: ZZ-type zinc finger; NLS: nuclear localisation signal; NES: nuclear export signal; TB: TRAF binding region; LIR: LC3 interacting region; KIR: Keap interacting region; UBA: ubiquitin associated domain). (B) Schematic overview of T4 polynucleotide kinase labelling assay (PNK). Cells are UV-crosslinked to establish a covalent bond between proteins and RNA at zero-distance. Subsequently, lysates are treated with RNase A and used for immunoprecipitation followed by radioactive labelling of RNA with T4 polynucleotide kinase. (C) PNK. Full-length (FL) p62 or different p62 truncations were expressed by transient transfection in HuH-7 p62 knockout cells and RNA binding capacity determined as described in B. (D) Quantification of PNK assays as in C. Adjusted P values are indicated according to one-way ANOVA with Tukey correction for multiple comparisons. (n=4). (E) Native immunoprecipitation of FLAG-HA-p62 full-length (FL) or truncation D from transfected HuH-7 p62 KO cells followed by quantitative RT-PCR of bound RNA. Indicated are adjusted p values from unpaired t-tests with Holm-Sidak correction for multiple comparisons (n=4). (vt1-1: vault RNA 1-1)

The N-terminal 172 amino acids of p62 carry the full RNA-binding capacity compared to the full-length (FL) p62 (Figure 1C and D). Additional deletion of the ZZ domain, leaving only the PB1 domain and adjacent C-terminal linker region of p62 (p62_1-122_; aa 1-122) significantly increases normalised RNA binding, suggesting a negative modulatory function of the ZZ domain on RNA binding. However, the PB1 domain alone (aa 1-102) or the ZZ domain with the linker (aa 102-172) display little if any RNA-binding capacity (Figure 1C and D). These data map the relevant RNA-binding interfaces to p62_1-122_. By contrast, the ZZ domain (aa 115-172) may play a regulatory role (Figure 1C and D; see below).

To quantitatively assess RNA binding under steady-state (i.e. non-crosslink) conditions and to investigate specific vault RNA1-1 binding, we performed native immunoprecipitation followed by RT-qPCR of co-purified RNAs (RIP). In line with the PNK assays, p62_1-122_ displays specific and maximal vault RNA1-1 binding compared to full-length p62 in this orthogonal assay (Figure 1E, Supplementary Figure 1). In contrast, p62 association with the other vault RNA paralogs is not significantly changed.

We conclude that p62_1-122_ -the PB1 domain with its adjacent C-terminal linker -accounts for specific vault RNA1-1 binding to p62. This result is unexpected in the light of the previously identified ZZ domain mutant R139A/K141A, which showed reduced RNA binding and implicated the ZZ domain as a critical region for the RNA interaction [19]. The data shown here shed new light on this mutant and suggest a regulatory role for the ZZ domain in RNA binding.

### Assignment of critical riboregulatory functions to Lys7 and Arg21

We next generated FLAG-HA tagged mutant constructs of p62 to i) assess, whether RNA binding depends on oligomerisation, and to ii) narrow down the RNA-binding interface in the context of full-length p62. To perturb PB1-dependent oligomerisation, we mutated the negatively charged OPCA motif (Figure 2A) introducing single (D69A), double (D69A/D71A) or triple amino acid (D69A/D71A/D73A) exchanges[11]. We assessed multimerization of the different constructs *in cellulo* by measuring the ratio of p62 complexes with slower migration in SDS-PAGE over the total p62 amounts in PNK assays. As expected, all mutants display decreased oligomerization compared to the wild-type control and to each other with increasing mutations (Figure 2B, Supplementary Figure 2B). Yet, their RNA-binding capacity was not significantly changed (Figure 2B, Supplementary Figure 2). In addition, we used maltose-binding protein (MBP)-tagging to generate recombinant p62 with low oligomerization potential [19,27,35] and tested *in vitro* binding of vault RNA1-1. This analysis confirms that the binding affinity of p62 is not affected by a double mutation of residues D69 and D73 (Supplementary Figure 2A).

**Figure 2:**
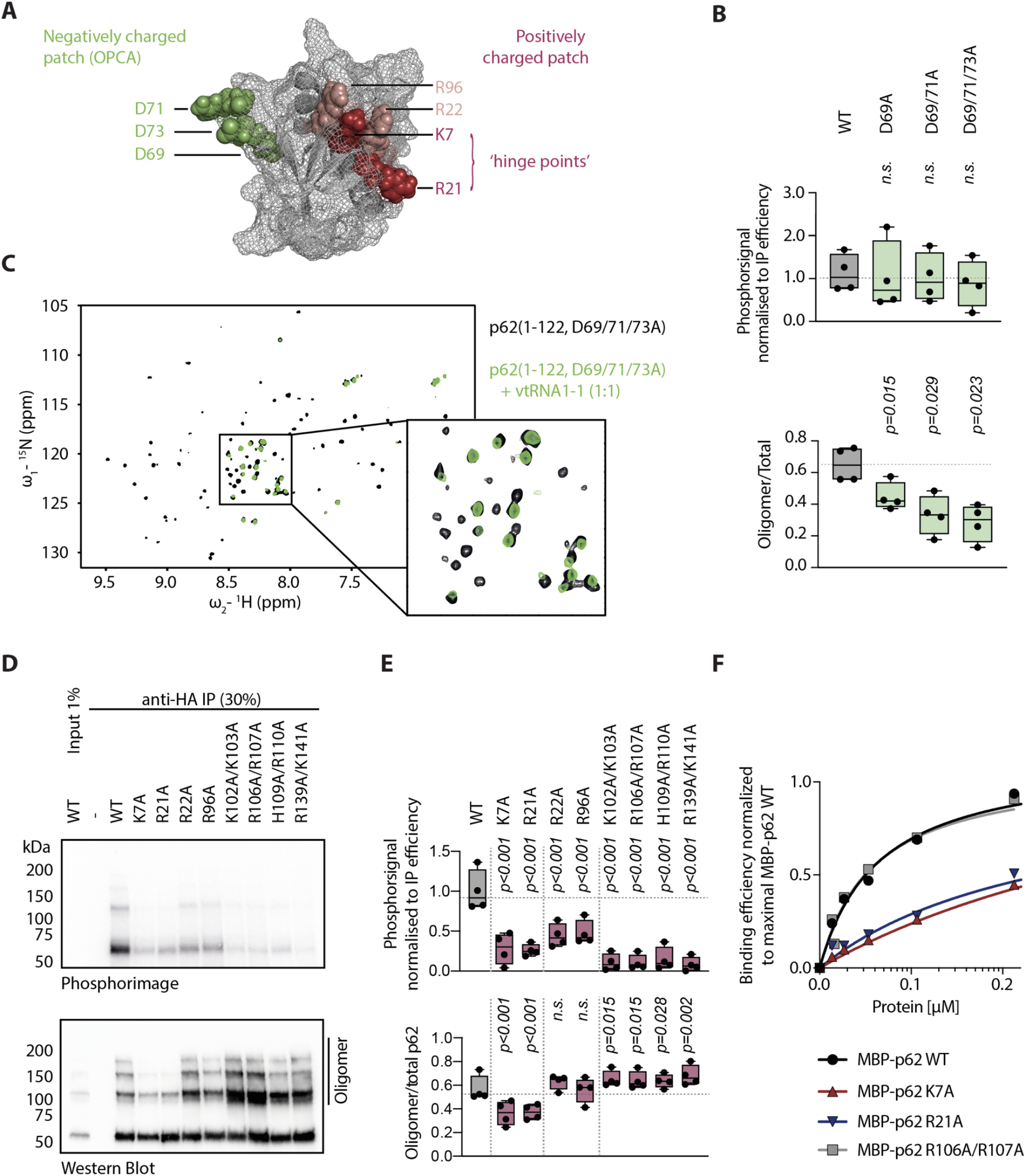
Lysine 7 and arginine 21 are hotspots for p62 riboregulation. (A) p62 PB1 domain structure (PDB ID: 2KKC). Negatively and positively charged residues are highlighted in green and red, respectively. (B) Quantification of radioactive signal and oligomerisation in PNK assays of full-length FLAG-HA-p62 oligomerisation mutants expressed in HuH-7 p62 KO cells. Significant differences from WT construct were assessed by RM one-way ANOVA with Benjamini and Hochberg correction for multiple comparisons. (n.s.: not significant; n=4) (C) ^1^H, ^15^N-HSQC NMR spectrum of p62(1-122, D69A/D71A/D73A) recorded with (green) and without (black) equimolar amounts of *in vitro* transcribed full-length vault RNA 1-1. Most peaks show dramatic intensity loss upon RNA addition. Zoom-in highlights chemical shift perturbations for the remaining sharp peaks in the central region. (D) Representative PNK assay of full-length FLAG-HA-p62 RNA binding mutants expressed in HuH-7 p62 KO cells. (E) Quantification of radioactive signal and oligomerisation in PNK assays of full-length FLAG-HA-p62 RNA binding mutants expressed in HuH-7 p62 KO cells. Significant differences from WT construct were assessed by RM one-way ANOVA with Benjamini and Hochberg correction for multiple comparisons. (n.s.: not significant; n=4) (F) Quantification of representative EMSA with 10 nM radioactively labelled vault RNA 1-1, 60 µM BSA, 150 nM bacterial tRNAs and increasing amounts of recombinantly expressed and purified MBP-p62 WT, MBP-p62 K7A, MBP-p62 K21A and MBP-p62 R106A/R107A.

Next, we performed ^1^H, ^15^N-HSQC NMR spectroscopy with the minimal construct p62_1-122_ which we forced into a monomeric form by the before-characterised triple mutation (D69A/D71A/D73A). We recorded spectra with and without equimolar amounts of *in vitro* transcribed full-length vault RNA1-1 to assess binding (Figure 2C). The spectra revealed strong signal loss for most peaks and multiple chemical shift perturbations for the remaining peaks upon addition of the RNA, indicating complex formation. Together, these findings show that the PB1 domain and adjacent C-terminal linker region of p62 are sufficient for direct and specific vault RNA1-1 binding and that oligomerisation is not required for this interaction.

To pinpoint the RNA-binding interface of p62, we mutated positively charged amino acids within the PB1 domain (Figure 2A) and adjacent linker region. This strategy yielded three classes of mutants (Figure 2D and E). First, PB1 domain mutants with decreased RNA binding and impaired oligomerization -namely p62 K7A and K21A. Second, those that display decreased RNA binding without significant changes in oligomerization, including R22A and R96A. And finally, the linker mutants K102A/K103A, R106A/R107A and H109A/R110A that show reduced RNA binding and increased multimerization. This latter group also includes the previously identified p62 ZZ domain mutant R139A/K141A[19], which we included here for reference.

Previous mutational studies showed that residues K7 and R21 are necessary for p62 multimerization, while R22 is not [11]. The PNK assay reflects this finding, confirming its utility as a tool to monitor p62 multimerization. Our data highlight residues K7 and R21 as particularly critical for riboregulation: they are required for both PB1 domain-mediated multimerization and RNA binding. As such, they appear to represent hinges for riboregulation.

Against that, R22 and R96 appear to extend the RNA-binding interface without making a strong contribution to multimerization. Mutations in the linker region or ZZ domain of p62 both resulted in reduced RNA binding and increased presence of multimers (Figure 2D and E). To distinguish whether in these cases the loss of RNA binding resulted in multimerization or conversely multimer formation displaced the RNA, we performed EMSAs with recombinant p62 that we forced into a low-oligomeric form by MBP-tagging [19,27,35]. We observed decreased vault RNA1-1 association for recombinant MBP-p62 K7A and R21A (Figure 2F, Supplementary Figure 2C and D), confirming the direct involvement of these residues in RNA binding. In contrast, the RNA-binding capacity was not impaired for the linker mutant MBP-p62 R106A/R107A (Figure 2F) or the ZZ domain mutant MBP-p62 R139A/K141A (Supplementary Figure 2E). These results favour the interpretation that mutations targeting the linker and ZZ domain foster multimeric forms of p62 *in cellulo* that exclude RNA from binding to the autophagic receptor.

### A central flexible loop of vault RNA 1-1 specifies p62 binding

The tertiary structure of vault RNA1-1 is presently unknown. Previous secondary structure analyses using RNase H probing were useful but limited in resolution [21,36]. We applied chemical structure probing in solution to identify relevant features of vault RNA1-1 at nucleotide resolution (Figure 3A and B). The low reactivity of the conserved ends of vault RNA1-1 towards the probing reagents and their high base complementarity suggest that the ends of vault RNA1-1 form a base-paired stem. This finding fits with previous thermodynamic models and is supported by the inter-species conservation of their base-pairing potential. In contrast, the central domain forms a highly reactive flexible loop region that was previously not anticipated.

**Figure 3:**
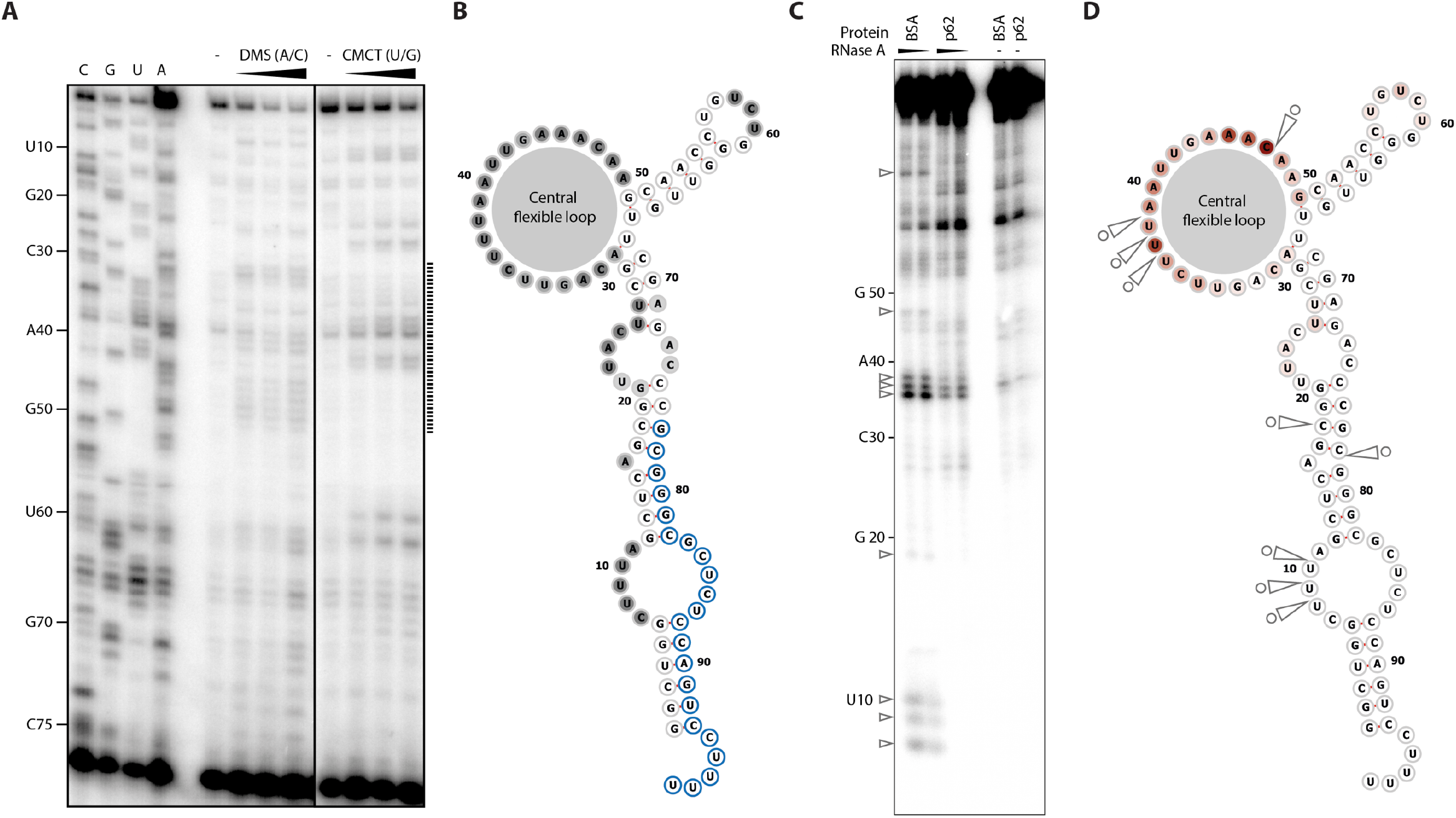
p62 interacts with a flexible loop in the central domain of vault RNA 1-1. (A) Chemical structure probing of *in vitro* transcribed vault RNA 1-1 in solution using Dimethyl sulphide (DMS) and 1-Cylcohexyl-(2-Morpholinoethyl)Carbodiimide metho-p-Toluene sulfonate (CMCT). The modified RNA is used for reverse transcription with a 5’-end labelled primer. The resulting cDNA is resolved on a denaturing urea-polyacrylamide gel. Sequencing lanes are depicted on the left side. Dotted line indicates highly reactive nucleotides within the central domain of vault RNA 1-1. (B) Secondary structure model of vault RNA 1-1 based on chemical structure probing. Residues with blue circle indicate the region of primer annealing for reverse transcription. The grey circle indicates the central flexible loop. (C) RNase A footprinting assay. *In vitro* transcribed vault RNA 1-1 was radiolabelled at the 5’-end, incubated with recombinant MBP-p62 or BSA and subjected to limited RNase A digest followed by precipitation. RNase protection patterns were resolved on a denaturing urea-polyacrylamide gel. Arrows indicate RNase A protected nucleotides. (D) Integration of secondary structure model with RNase protection sites (arrows) from C and mean crosslink site values of vault RNA1-1 from p62-iCLIP analysis (red shading indicates increased crosslinking with p62). (The iCLIP data presented in this figure were adapted from [19])

Next, we conducted RNase A footprinting to identify vault RNA1-1 residues that are protected by p62 binding (Figure 3C and D, grey arrows). Several protected regions emerge in the presence of p62. These include single-stranded bulges within the stem and, prominently, the central flexible loop. Of note, the protected nucleotides in the central loop completely match the crosslinked nucleotides previously identified by iCLIP of p62 (Figure 3D, red shading [19]). When compared with the other three human vault RNA paralogs, the central loop of vault RNA1-1 differs considerably in length and sequence (Supplementary Figure 3). Overall, our data identify nucleotides within the central loop of vault RNA 1-1 as specific p62 contact points.

### The central flexible loop of vault RNA1-1 determines p62 riboregulation *in cellulo*

To validate the importance of the central flexible loop of vault RNA 1-1 functionally for riboregulation, we generated a HuH-7 Flp-IN vault RNA1-1 KO cell line with a single FRT integration site. This KO cell line thus allowed stable re-integration of vault RNA1-1 variants expressed from the endogenous promoter (Supplementary Figure 4). We compared re-integrated wild-type vault RNA1-1 with two mutants that target either the central loop triplet U36/U37/U38 alone (M1) or in combination with the loop residues A46/C47 (M2; Figure 4A). The expression levels of all three re-integrated vault RNA1-1 variants are similar to each other and correspond to approximately 50% of the vault RNA1-1 levels in the parental wild type HuH-7 Flp-IN cell line (Figure 4B and C). This expression level is in line with the single FRT integration site and, expectedly, also reflected in a reduced vault RNA1-1 association with p62 as measured by native IP of p62 followed by RT-qPCR quantification (Figure 4D, Supplementary figure 4E). As expected, the central domain mutants M1 and M2 tend to show reduced p62 binding *in cellulo* compared to the wild-type vault RNA1-1 (Figure 4D, Supplementary figure 4E). This conclusion is further supported *in vitro* by EMSAs, where the mutations in the central loop of vault RNA1-1 compete less well for p62 binding than the wt counterpart (Supplementary Figure 5A). This assay also shows that the central loop region of vtRNA1-1 alone is not sufficient for p62 binding *in vitro* (Supplementary Figure 5A) and reveals the importance of the 3D structure of vtRNA1-1 for p62 binding.

Finally, we tested the loop mutations for effects on p62 riboregulation. As established before [19], we stimulated autophagy in the reconstituted cells with the synthetic, p62-specific ZZ-domain ligand XIE62-1004-A [6] and assessed the LC3II/LC3I ratio as a measure of autophagic flux by Western blotting. Consistent with our earlier data, the autophagic flux is significantly greater in cells depleted of vault RNA1-1 compared to the respective CRISPR/Cas9 control cell line upon treatment, as evidenced by the increased ratio of LC3BII/LC3BI (Figure 4E and F). Re-integration of wild-type vault RNA1-1 rescues this phenotype significantly, whereas the central loop mutants fail to do so (Figure 4E and F). Thus, the central loop of vault RNA1-1 is also required for efficient riboregulation of p62-mediated autophagy *in cellulo*.

**Figure 4:**
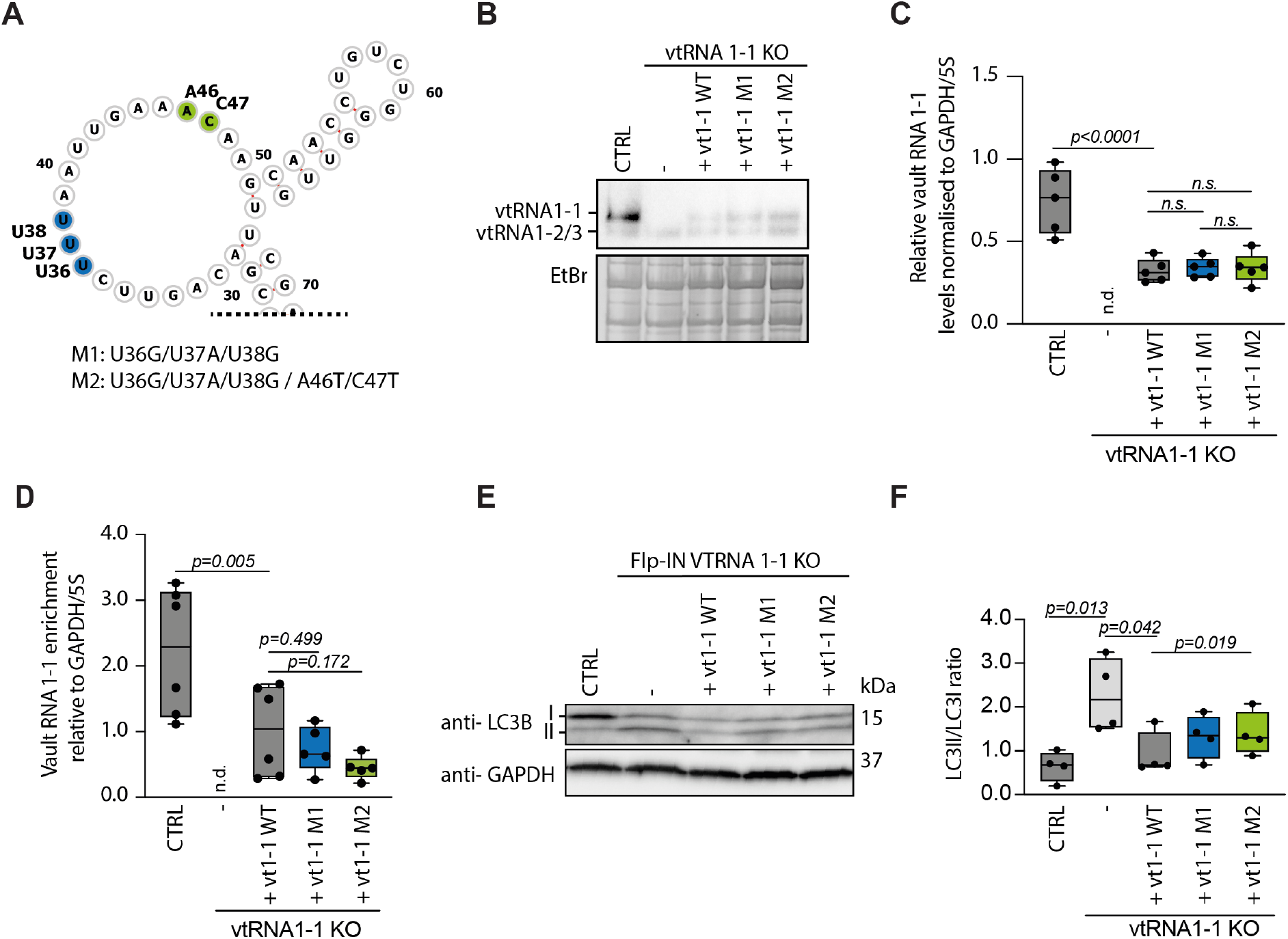
The flexible loop region is a key determinant of p62 riboregulation in cellulo. (A) Schematic representation of the flexible loop region. Site of mutations are indicated by blue and green shading. (B) Northern blot analysis of HuH-7 Flp-IN cell lines that were used to stably express vault RNA 1-1 and mutants thereof in a vault RNA 1-1 KO background (please see Supplementary Figure 4A). Vault RNA 1-1, 1-2 and 1-3 were detected with a vault RNA specific, mutation-independent probe. (C) RT-qPCR. Expression of vault RNA 1-1 and mutants thereof was detected with a specific, mutation-independent primer pair. Individual p values are indicated according to one-way ANOVA with Benjamini and Hochberg correction for multiple comparisons. (n=5) (D) Native p62 RIP followed by RT-qPCR. Vault RNA 1-1 and mutants thereof were detected with specific, mutation-independent primer pair. Individual p values are indicated according to one-way ANOVA with Benjamini and Hochberg correction for multiple comparisons. (n≥5). (E) Western blot analysis of LC3B ratio. The cell lines were treated with 10 µM XIE62-1004-A for 4 h to induce p62-specific autophagy. Lysates were analysed by Western blot. (F) Quantification of LC3BII/I ratio in Western blot analysis as performed in E). Individual p values are indicated according to RM one-way ANOVA with Benjamini and Hochberg correction for multiple comparisons. Significance was assessed by paired Student’s t-test. (n=4).

## DISCUSSION

Earlier work unveiled riboregulation as a new modality of control for autophagy: to limit autophagic flux, p62 oligomerization is inhibited by direct vtRNA1-1 binding [18,19]. This discovery raised the question of how vtRNA1-1 interferes with oligomerization and how specificity is achieved in comparison to the other three human vault RNA orthologs and other tRNA-like Pol III transcripts. The work presented here addresses these mechanistic questions and uncovers critical features that determine binding and specificity of the p62-vtRNA1-1 riboregulatory pair. It reveals molecular details of how vtRNA1-1 modulates p62 function by interference with PB1 domain-mediated oligomerisation.

We show that the PB1 domain and adjacent linker region (p62_1-122_) are both necessary and sufficient for maximal specific vtRNA1-1 binding. Mutational analysis pinpoints K7 and R21 as critical amino acids that are necessary for RNA binding (Figure 2) in addition to their previously known role in p62 oligomerisation [11]. This result identifies these two residues as hinge points for riboregulation, through which vtRNA1-1 inhibits p62 oligomerisation and consequently autophagy.

Our initial report identified two mutants in the ZZ domain (K141A and R139A/K141A) that showed strongly diminished RNA binding and increased p62 oligomerization ([19]; see also Figure 2D and E). Interestingly, the three new linker domain mutants K102A/K103A, R106A/R107A and H109A/R110A display the same phenotype (Figure 2E and F). Importantly, the ZZ domain had previously been shown to bind arginylated substrates (N-degrons) [6,17], a process that triggers p62 oligomerisation and thereby initiates autophagic clearance. In this context, the linker region (p62_100-113_) was shown to exert negative autoregulation through direct interaction with the ZZ domain [17]. In the light of our new data (Figure 2), we hypothesize that the above mutations within the linker region or the ZZ domain, respectively, render this autoinhibitory mechanism dysfunctional, resulting in constitutive activation of the clearance pathway and increased p62 oligomerization. Of note, the auto-regulatory linker overlaps exactly with all PB1 domain mutants that show a significant increase in multimerization. In further support of this model, R139 has been shown to directly interact with the inhibitory linker and stabilise the autoregulatory interaction [17]. The exclusion of RNA from p62 oligomers that are formed upon activation of ZZ domain-mediated autophagic clearance would ensure efficient aggregate clearance even in situations when cellular levels of the vault RNA1-1 riboregulator are high, including for example viral infections[21,37–39].

As proposed before, our refined model suggests that vault RNA1-1 primarily inhibits the process of oligomerisation, but may not be able to disrupt p62 oligomers that are formed when cargo binds to the ZZ domain. Integrating all available experimental evidence, we suggest the following mechanistic model for riboregulation of p62 by vtRNA1-1: under physiological, nutrient-replete conditions, vtRNA1-1 inhibits p62 oligomerisation by binding the critical hinge points K7 and R21. When starvation reduces cellular vtRNA1-1 levels, p62 oligomerisation and autophagy are facilitated (Supplementary Figure 5B; [19]). By contrast, cargo binding to the ZZ domain and linker region during proteotoxic stress [17,40] triggers ‘sequestosome’ formation and cargo clearance even when vtRNA1-1 levels are high, excluding vtRNA1-1 sterically in a dominant fashion (Supplementary Figure 5B).

To also decipher the riboregulatory interface(s) on vtRNA1-1, we defined the nucleotides required for p62 binding (Figure 3-4). We discovered that a central loop region of vtRNA1-1 is necessary for this interaction and for specific riboregulation of p62 function in autophagy. Interestingly, this region of vtRNA1-1, especially nucleotides 45-50, had previously been shown to mediate apoptosis resistance in Hela cells [37,41]. While the molecular details of this phenotype remain unclear, our finding suggests the modulation of autophagy-dependent apoptosis via p62 as a possibility deserving of further exploration.

The p62-vtRNA1-1 binding interface may well be targeted by regulatory modifications that potentially influence riboregulation. For example, the hinge point K7 in the PB1 domain of p62 has previously been found to be ubiquitinated by TRIM21 [16]. Like vtRNA1-1 binding, this ubiquitination prevents p62 oligomerisation, but also facilitates proteasomal degradation of p62. By contrast, vtRNA1-1 riboregulation may represent a more dynamic and reversible form of autophagy modulation. It will be interesting to examine this critical riboregulatory interface for other protein or RNA modifications that could represent additional layers of regulation.

Our new insights on the binding interface could potentially help to modulate p62-specific autophagy and aggregate clearance by reversibly changing the expression levels of the riboregulatory vault RNA 1-1, for example through RNA-interference pathways. This might be of special benefit in instances, where the cargo load exceeds the capacity of cellular degradation mechanisms as it is the case in neurodegenerative diseases or cancer.

In addition to its function as an autophagy receptor, p62 provides a platform for key cellular signalling pathways [42–45]. While we have not detected changes in mTOR signalling upon depletion of vtRNA1-1 in HuH-7 cells [19], it will be interesting to explore globally whether RNA1-1 depletion or RNA-binding deficient p62 mutants affect other cellular signalling pathways.

With hundreds of newly identified RNA-binding proteins [46] riboregulation could be involved in the modulation of many key cellular processes beyond autophagy. This report offers a paradigm for the molecular details underlying riboregulation. These may also serve in the design of new strategies to modulate this key cellular process involved in ageing, cancer and neurodegenerative diseases in the future.

## ACKNOWLEDGEMENT

We thank Dmytro Dziuba for the synthesis of the XIE62-1004-A compound. Special thanks to Ina Huppertz for fruitful discussions and critical reading of the manuscript.

## AUTHOR CONTRIBUTIONS

M.W.H. and M.B. designed the project and wrote the manuscript with input from all authors. M.B. performed and analysed most experiments with input from all authors. R.H. and M.W.H. co-supervised the study. F.B. was essential to the setup and analysis of RNA structure probing and footprinting. N.D and K.H. performed protein purifications for NMR. K.H. and J.H. performed and analysed the NMR experiments.

## REFERENCES

1. Bjørkøy G, Lamark T, Brech A et al. p62/SQSTM1 forms protein aggregates degraded by autophagy and has a protective effect on huntingtin-induced cell death. J Cell Biol 2005;171:603–14.

2. Dikic I. Proteasomal and Autophagic Degradation Systems. Annual Review of Biochemistry 2017;86:193–224.

3. Lim J, Lachenmayer ML, Wu S et al. Proteotoxic Stress Induces Phosphorylation of p62/SQSTM1 by ULK1 to Regulate Selective Autophagic Clearance of Protein Aggregates. PLOS Genetics 2015;11:e1004987.

4. Jakobi AJ, Huber ST, Mortensen SA et al. Structural basis of p62/SQSTM1 helical filaments and their role in cellular cargo uptake. Nat Commun 2020;11:1–15.

5. Zaffagnini G, Savova A, Danieli A et al. p62 filaments capture and present ubiquitinated cargos for autophagy. EMBO J 2018;37, DOI: 10.15252/embj.201798308.

6. Cha-Molstad H, Yu JE, Feng Z et al. p62/SQSTM1/Sequestosome-1 is an N-recognin of the N-end rule pathway which modulates autophagosome biogenesis. Nature Communications 2017;8:102.

7. Pankiv S, Clausen TH, Lamark T et al. p62/SQSTM1 Binds Directly to Atg8/LC3 to Facilitate Degradation of Ubiquitinated Protein Aggregates by Autophagy. J Biol Chem 2007;282:24131–45.

8. Wurzer B, Zaffagnini G, Fracchiolla D et al. Oligomerization of p62 allows for selection of ubiquitinated cargo and isolation membrane during selective autophagy. eLife Sciences 2015;4:e08941.

9. Ciuffa R, Lamark T, Tarafder AK et al. The Selective Autophagy Receptor p62 Forms a Flexible Filamentous Helical Scaffold. Cell Reports 2015;11:748–58.

10. Ito T, Matsui Y, Ago T et al. Novel modular domain PB1 recognizes PC motif to mediate functional protein–protein interactions. EMBO J 2001;20:3938–46.

11. Lamark T, Perander M, Outzen H et al. Interaction Codes within the Family of Mammalian Phox and Bem1p Domain-containing Proteins. J Biol Chem 2003;278:34568–81.

12. Itakura E, Mizushima N. p62 targeting to the autophagosome formation site requires self-oligomerization but not LC3 binding. J Cell Biol 2011;192:17–27.

13. Sun D, Wu R, Zheng J et al. Polyubiquitin chain-induced p62 phase separation drives autophagic cargo segregation. Cell Res 2018;28:405–15.

14. Carroll B, Otten EG, Manni D et al. Oxidation of SQSTM1/p62 mediates the link between redox state and protein homeostasis. Nature Communications 2018;9:256.

15. Christian F, Krause E, Houslay MD et al. PKA phosphorylation of p62/SQSTM1 regulates PB1 domain interaction partner binding. Biochimica et Biophysica Acta (BBA) - Molecular Cell Research 2014;1843:2765–74.

16. Pan J-A, Sun Y, Jiang Y-P et al. TRIM21 ubiquitylates SQSTM1/p62 and suppresses protein sequestration to regulate redox homeostasis. Mol Cell 2016;61:720–33.

17. Zhang Y, Mun SR, Linares JF et al. ZZ-dependent regulation of p62/SQSTM1 in autophagy. Nat Commun 2018;9, DOI: 10.1038/s41467-018-06878-8.

18. Büscher M, Horos R, Hentze MW. ‘High vault-age’: non-coding RNA control of autophagy. Open Biology 2020;10:190307.

19. Horos R, Büscher M, Kleinendorst R et al. The Small Non-coding Vault RNA1-1 Acts as a Riboregulator of Autophagy. Cell 2019;176:1054-1067.e12.

20. Kedersha NL, Rome LH. Isolation and characterization of a novel ribonucleoprotein particle: large structures contain a single species of small RNA. J Cell Biol 1986;103:699–709.

21. Nandy C, Mrázek J, Stoiber H et al. Epstein–Barr Virus-Induced Expression of a Novel Human Vault RNA. Journal of Molecular Biology 2009;388:776–84.

22. Stadler PF, Chen JJ-L, Hackermüller J et al. Evolution of vault RNAs. Mol Biol Evol 2009;26:1975–91.

23. Beckmann BM, Horos R, Fischer B et al. The RNA-binding proteomes from yeast to man harbour conserved enigmRBPs. Nat Commun 2015;6:1–9.

24. van den Ent F, Löwe J. RF cloning: A restriction-free method for inserting target genes into plasmids. Journal of Biochemical and Biophysical Methods 2006;67:67–74.

25. Cong L, Ran FA, Cox D et al. Multiplex genome engineering using CRISPR/Cas systems. Science 2013;339:819–23.

26. Ran FA, Hsu PD, Wright J et al. Genome engineering using the CRISPR-Cas9 system. Nat Protoc 2013;8:2281–308.

27. Tarafder AK, Guesdon A, Kuhm T et al. Recombinant Expression, Purification, and Assembly of p62 Filaments. In: Ktistakis N, Florey O (eds.). Autophagy: Methods and Protocols. New York, NY: Springer, 2019, 3–15.

28. Simon B, Köstler H. Improving the sensitivity of FT-NMR spectroscopy by apodization weighted sampling. J Biomol NMR 2019;73:155–65.

29. Delaglio F, Grzesiek S, Vuister GW et al. NMRPipe: A multidimensional spectral processing system based on UNIX pipes. J Biomol NMR 1995;6:277–93.

30. Lee W, Tonelli M, Markley JL. NMRFAM-SPARKY: enhanced software for biomolecular NMR spectroscopy. Bioinformatics 2015;31:1325–7.

31. Schindelin J, Arganda-Carreras I, Frise E et al. Fiji - an Open Source platform for biological image analysis. Nat Methods 2012;9, DOI: 10.1038/nmeth.2019.

32. Greenberg JR. Ultraviolet light-induced crosslinking of mRNA to proteins. Nucleic Acids Res 1979;6:715–32.

33. Pashev IG, Dimitrov SI, Angelov D. Crosslinking proteins to nucleic acids by ultraviolet laser irradiation. Trends in Biochemical Sciences 1991;16:323–6.

34. Suchanek M, Radzikowska A, Thiele C. Photo-leucine and photo-methionine allow identification of protein-protein interactions in living cells. Nature Methods 2005;2:261–8.

35. Reuten R, Nikodemus D, Oliveira MB et al. Maltose-Binding Protein (MBP), a Secretion-Enhancing Tag for Mammalian Protein Expression Systems. PLOS ONE 2016;11:e0152386.

36. Poderycki MJ, Rome LH, Harrington L et al. The p80 homology region of TEP1 is sufficient for its association with the telomerase and vault RNAs, and the vault particle. Nucleic Acids Res 2005;33:893–902.

37. Amort M, Nachbauer B, Tuzlak S et al. Expression of the vault RNA protects cells from undergoing apoptosis. Nature Communications 2015;6:7030.

38. Li F, Chen Y, Zhang Z et al. Robust expression of vault RNAs induced by influenza A virus plays a critical role in suppression of PKR-mediated innate immunity. Nucleic Acids Res 2015;43:10321–37.

39. Mrázek J, Kreutmayer SB, Grässer FA et al. Subtractive hybridization identifies novel differentially expressed ncRNA species in EBV-infected human B cells. Nucleic Acids Res 2007;35:e73.

40. Cha-Molstad H, Lee SH, Kim JG et al. Regulation of autophagic proteolysis by the N-recognin SQSTM1/p62 of the N-end rule pathway. Autophagy 2018;14:359–61.

41. Bracher L, Ferro I, Pulido-Quetglas C et al. Human vtRNA1-1 Levels Modulate Signaling Pathways and Regulate Apoptosis in Human Cancer Cells. Biomolecules 2020;10:614.

42. Duran A, Amanchy R, Linares JF et al. p62 is a key regulator of nutrient sensing in the mTORC1 pathway. Mol Cell 2011;44:134–46.

43. Katsuragi Y, Ichimura Y, Komatsu M. p62/SQSTM1 functions as a signaling hub and an autophagy adaptor. The FEBS Journal 2015;282:4672–8.

44. Lee SJ, Pfluger PT, Kim JY et al. A functional role for the p62–ERK1 axis in the control of energy homeostasis and adipogenesis. EMBO Rep 2010;11:226–32.

45. Moscat J, Karin M, Diaz-Meco MT. p62 in Cancer: Signaling Adaptor Beyond Autophagy. Cell 2016;167:606–9.

46. Gebauer F, Schwarzl T, Valcárcel J et al. RNA-binding proteins in human genetic disease. Nature Reviews Genetics 2020:1–14.

47. Raden M, Ali SM, Alkhnbashi OS et al. Freiburg RNA tools: a central online resource for RNA-focused research and teaching. Nucleic Acids Res 2018;46:W25–9.

48. Will S, Reiche K, Hofacker IL et al. Inferring noncoding RNA families and classes by means of genome-scale structure-based clustering. PLoS Comput Biol 2007;3:e65–e65.

49. Will S, Joshi T, Hofacker IL et al. LocARNA-P: accurate boundary prediction and improved detection of structural RNAs. RNA 2012;18:900–14.

