## Supplementary Material for "Mechanism of riboregulation of p62 protein oligomerisation by vault RNA1-1 in selective autophagy"

**Supplementary Figure 1:**

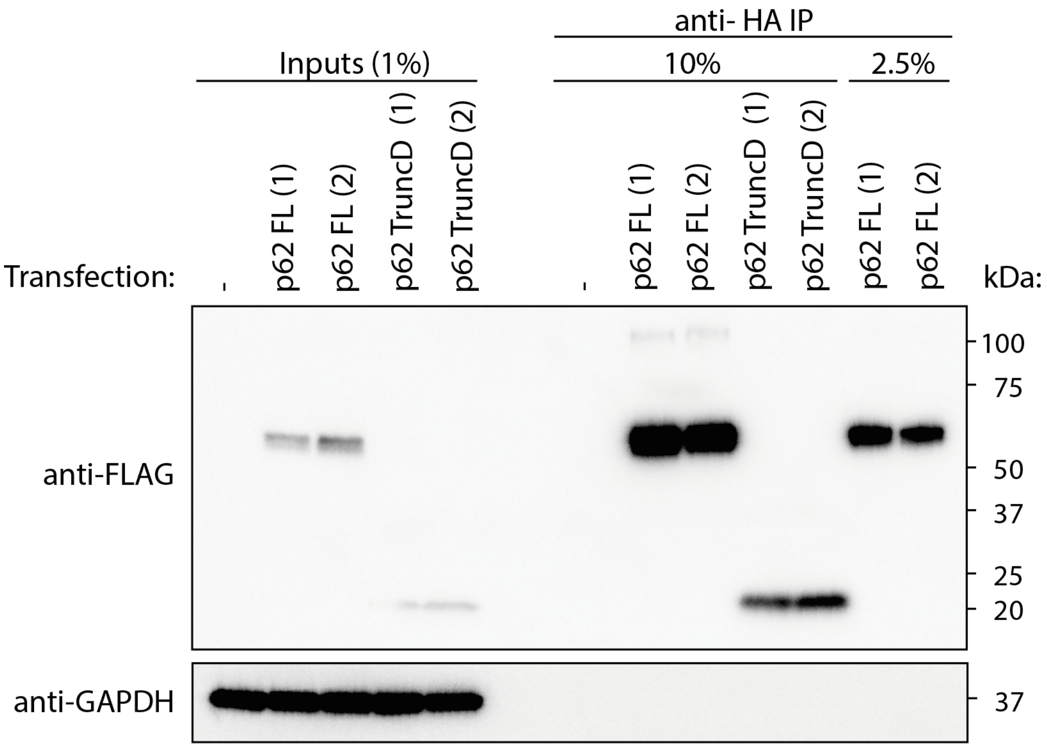


Native co-immunoprecipitation of FLAG-HA-p62 WT or truncation D expressed in HuH-7 p62 KO cells followed by quantitative RT-PCR of bound RNA. Representative Western blot analysis of p62 immunoprecipitation, including the normalization of eluates according to the protein content for subsequent RNA extraction.

**Supplementary Figure 2:**

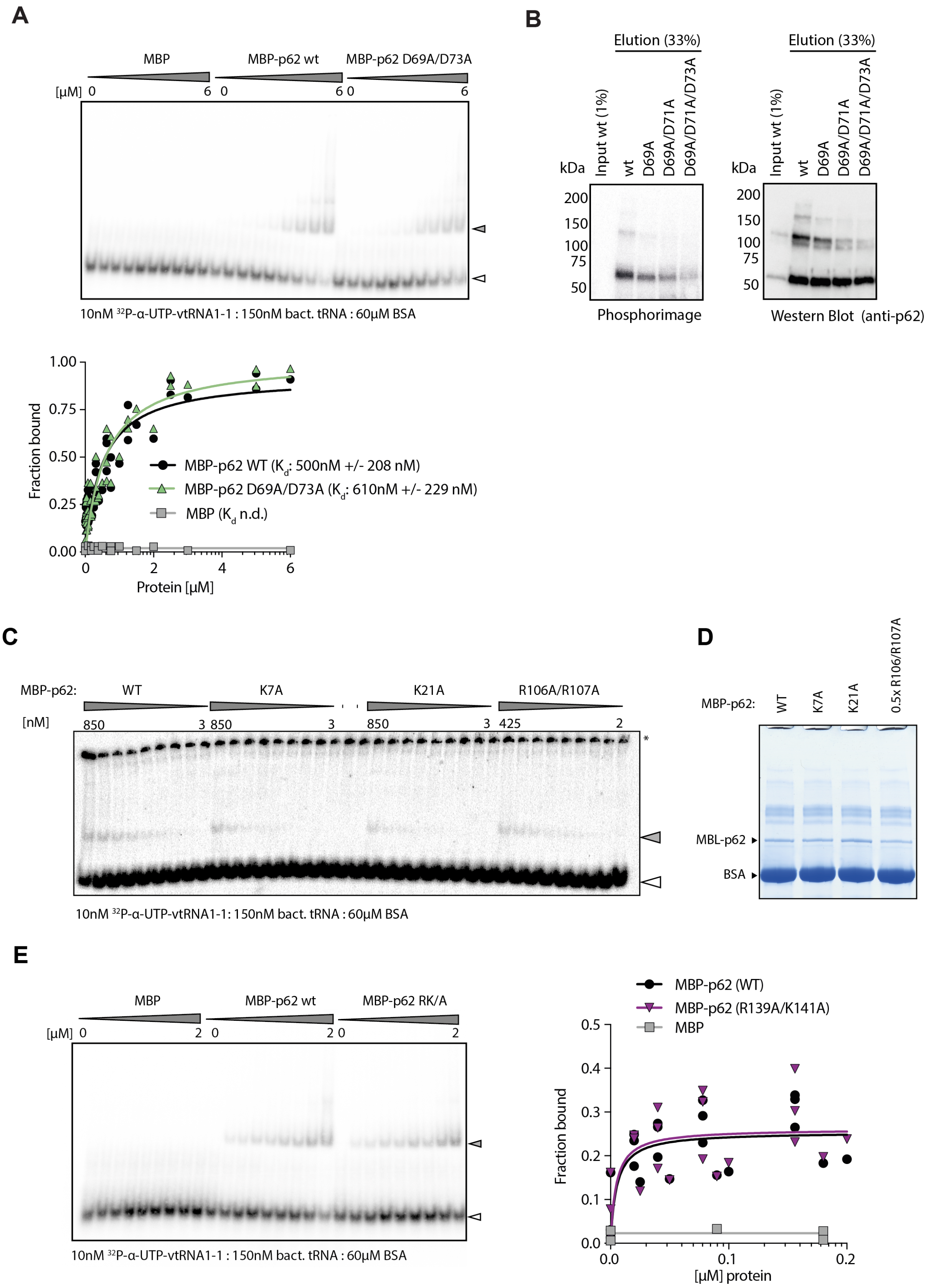


(A) Representative EMSA and quantification. Radioactively labelled vault RNA 1-1 and increasing amounts of recombinantly expressed and purified MBP (n=2), MBP-p62 WT (n=5) or MBP-p62 (D69A/D73A) (n=5) in the presence of an unspecific competitor.

(B) Representative Polynucleotide kinase labelling assay (PNK) of FLAG-HA-p62 mutants in HuH-7 p62 KO cells.
(C) Representative EMSA with 10 nM radioactively labelled vault RNA 1-1, 60 µM BSA, 150 nM bacterial tRNAs and increasing amounts of recombinantly expressed and purified MBP-p62 WT, MBP-p62 K7A, MBP-p62 K21A and MBP-p62 R106A/R107A. A white arrow indicates free radioactively labelled probe, a grey arrow indicates RNA-protein complex, * indicates the wells.
(D) SDS-PAGE followed by InstantBlue staining of EMSA protein reaction from B.
(E) Representative EMSA and quantification. Radioactively labelled vault RNA 1-1 and increasing amounts of recombinantly expressed and purified MBP-p62 WT (n=5), MBP-p62 R139A/K141A (n=5) or MBP (n=2) in the presence of unspecific competitor (data also partly shown in Figure 2D).

**Supplementary Figure 3:**

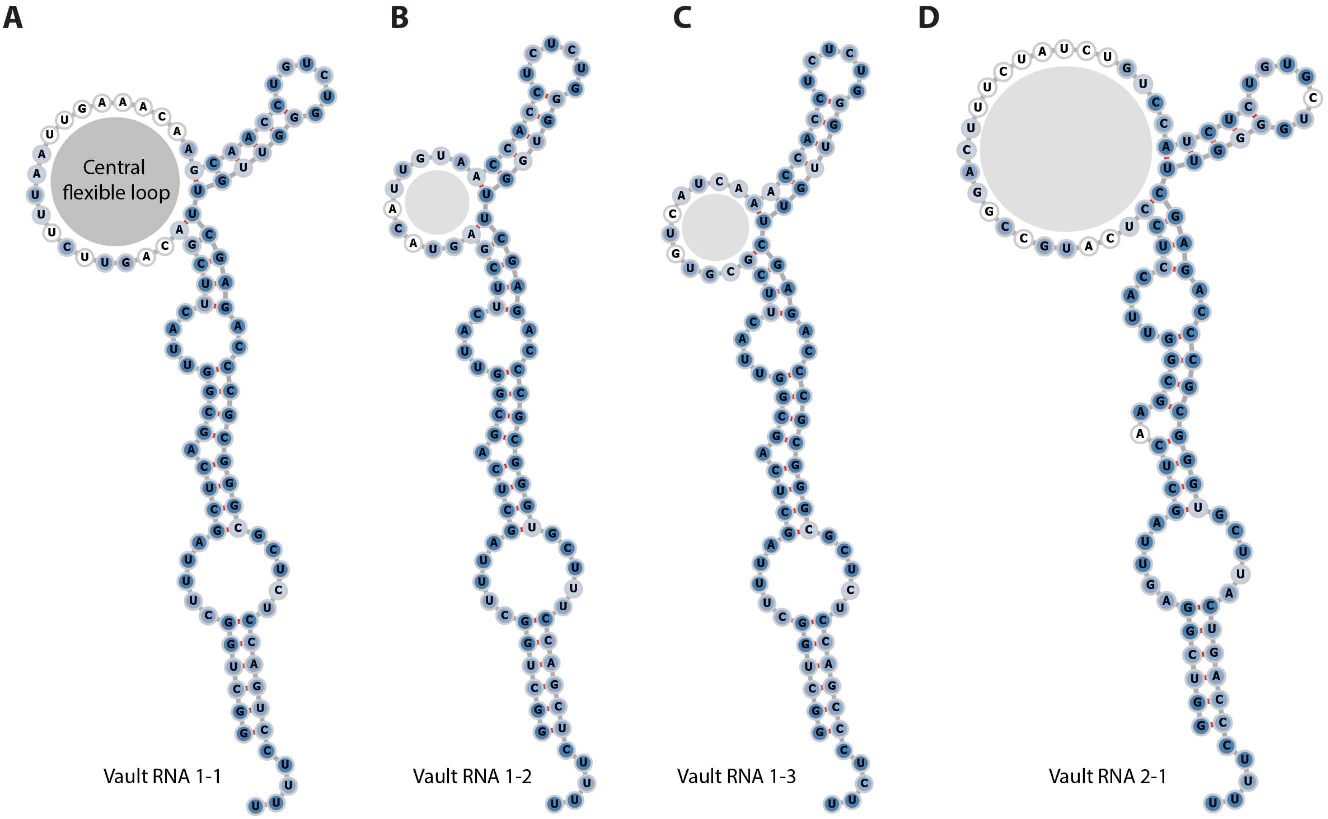


(A) Integration of secondary structure model with vault RNA paralogue conservation. The intensity of blue shading represents higher paralogue conservation as assessed by *LocARNA (http://rna.informatik.uni-freiburg.de, v. (4.5.8);* [47–49].
(B) – (D) Secondary structure models for the other human vault RNAs based on chemical probing of vault RNA 1-1 and paralogue conservation as in A.

**Supplementary Figure 4:**

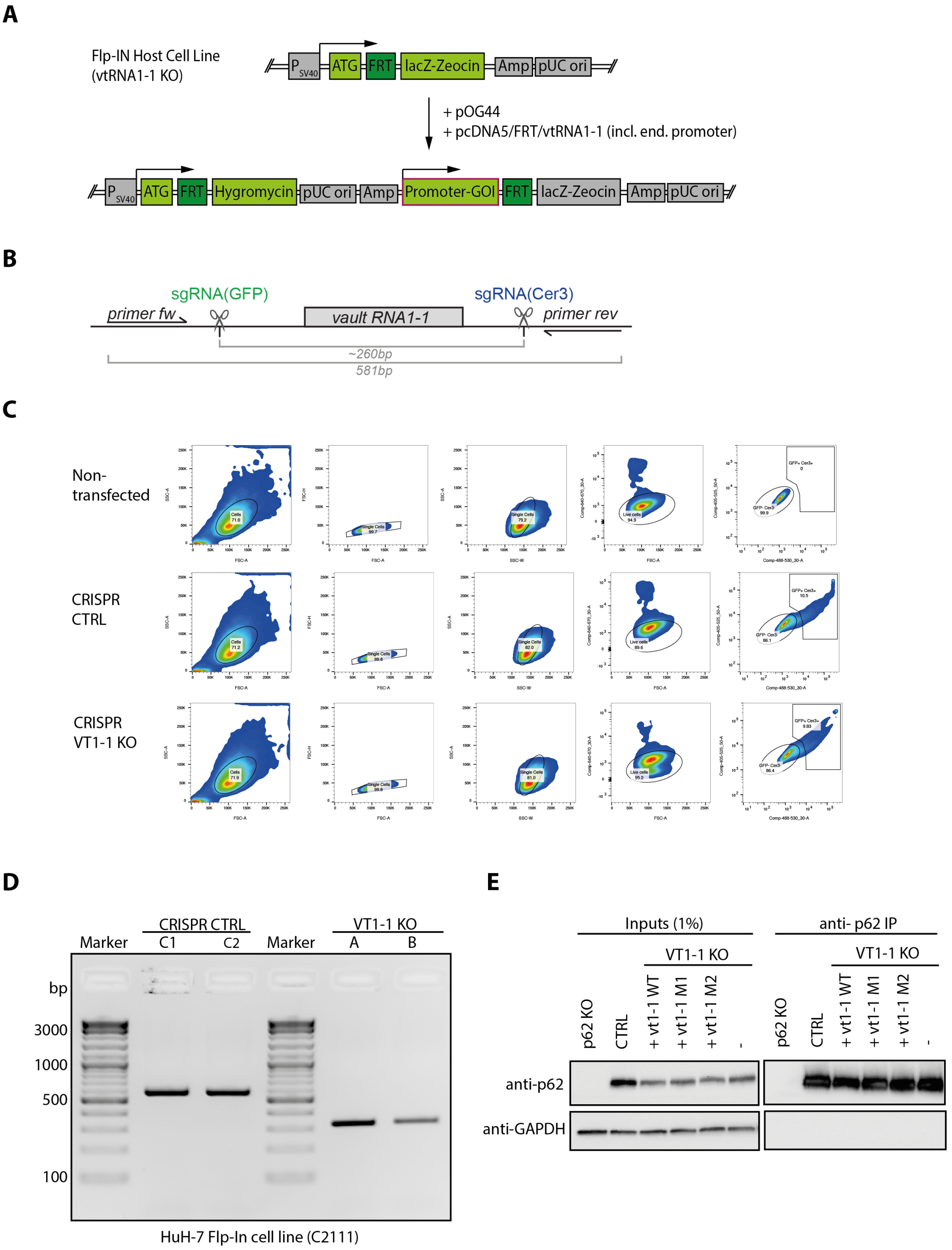


(A) Schematic overview of Flp-IN cell line for the reintegration of vault RNA 1-1 and mutants thereof. A gene of interest can be introduced through Flp-FRT recombination by co-transfection of an FRT site containing plasmid with the respective recombinase (pOG44). Upon successful integration, the cells gain hygromycin resistance and loose zeocin resistance.

(B) Schematic representation of the vault RNA 1-1 locus with localisation of single-guide RNAs (sgRNAs) and primers for PCR analysis.

(C) Single-cell FACS sorting of double-positive HuH-7 Flp IN cells that express both single guide RNAs targeting vault RNA 1-1 and CRISPR/Cas9.

(D) PCR analysis of genomic vault RNA1-1 loci of two single-cell derived HuH-7 Flp-IN CRISPR/Cas9 Control cell lines and vault RNA 1-1 KO cell lines.

(E) Native co-immunoprecipitation of p62 in HuH-7 FlpIN cells followed by quantitative RT-PCR of bound RNA. Representative Western blot analysis of p62 immunoprecipitation.

### Supplementary Figure 5:

**A**

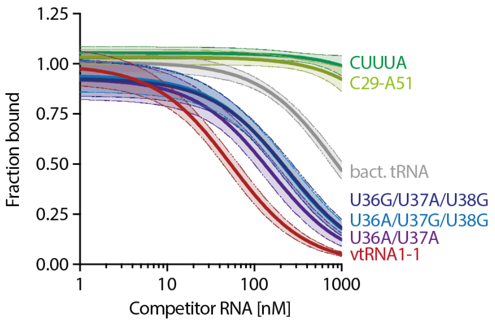


**B**

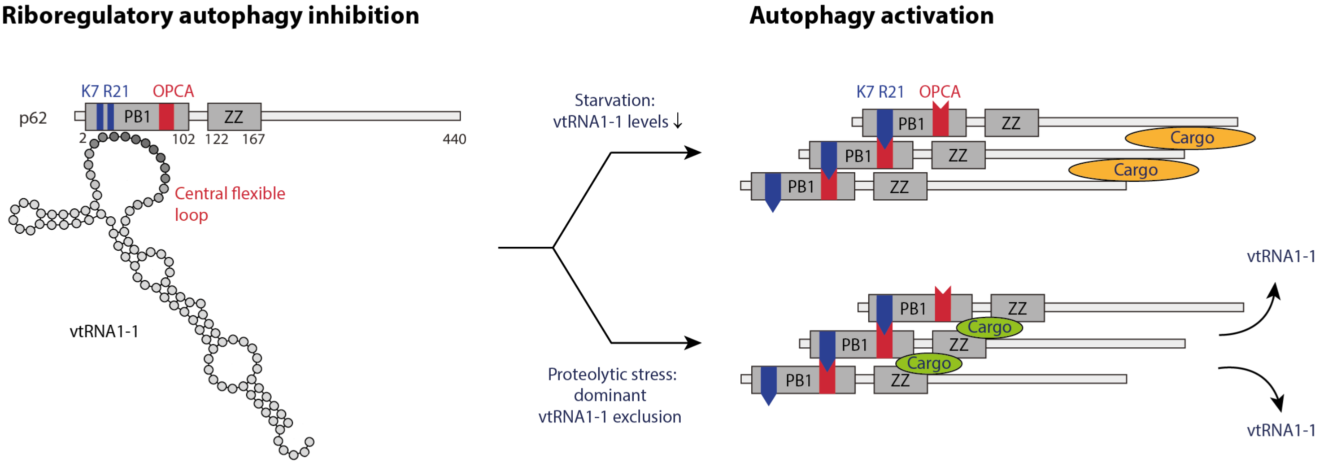


(A) Quantification of competitive EMSAs with vtRNA1-1 WT or full-length mutants, bacterial tRNAs, or synthetic RNA oligos representing the linear flexible loop region or the motif CUUUA (n≥2). Shading indicates 95% confidence interval. The analysis was performed with Prism 8.

(B) Summary model. Under physiological, nutrient-replete conditions, vtRNA1-1 inhibits p62 oligomerisation by binding the critical hinge points K7 and R21. When starvation reduces cellular vtRNA1-1 levels, p62 oligomerisation and autophagy are facilitated[19]. By contrast, cargo binding to the ZZ domain and linker region during proteotoxic stress[17,40] triggers 'sequestosome' formation and cargo clearance even when vtRNA1-1 levels are high, excluding vtRNA1-1 sterically in a dominant fashion. (blue: positively charged surface patch including K7 and R21, red: negatively charged OPCA motif; PB1: Phox and Bem1; ZZ: ZZ-type zinc finger; orange: cargo targeted for degradation via the UBA binding domain; green: cargo that binds the ZZ domain and linker region upon proteotoxic stress.

**Supplementary Table 1:**

| | | |
|---|---|---|
| **Antibodies** | | |
| Anti-p62 rabbit pAb | MBL | Cat#: PM045 |
| Anti-GAPDH rabbit pAB | Sigma-Aldrich | Cat#: G9545;<br>RRID: AB_796208 |
| Anti-FLAG mouse mAb | Sigma-Aldrich | Cat#: F1804-50UG |
| Anti-HA magnetic beads | Thermo Scientific | Cat#: 88836 |
| **Bacterial Strains** | | |
| *E.coli* TOP10 | Thermo Scientific | Cat#: C404010 |
| *E.coli* BL21(DE3) CodonPlus-RIL | Agilent | Cat#: 230240 |
| *E.coli* BL21 Rosetta$^{TM}$ 2 (DE3) | Sigma-Aldrich | Cat#: 71400 |
| **Chemicals, Peptides, and Recombinant Proteins** | | |
| InstantBlue Protein stain | Expedeon | Cat#: ISB1L |
| Precision Plus Protein Dual Color | Biorad | Cat#: 1610374 |
| XIE62-1004-A | [19] | Synthesized by D. Dziuba |
| AMV Reverse Transcriptase | Promega | Cat#: M5101 |
| Benzonase (100U/ml) | Merck Millipore | Cat#: 71206 |
| cOmplete, EDTA free | Sigma-Aldrich | Cat#: 11873580001 |
| FastAP alkalische phosphatase | Thermo Scientific | Cat#: EF0651 |
| FastDigest BbsI/BpiI | Thermo Scientific | Cat#: FD1014 |
| Phusion HF DNA polymerase | NEB | Cat#: M0530S |
| Quickligase | NEB | Cat#: M22000S |
| RNaseA | Sigma-Aldrich | Cat#: R5503 |
| T4 Polynucleotid kinase (PNK) | NEB | Cat#: M0201L |
| Turbo Dnase | Thermo Fisher | Cat#: AM2238 |
| **Critical Commercial Assays** | | |
| ChromaSpin + TE-10 columns | Takara | Cat#: 636066 |
| Fast SYBR Green Master Mix | Thermo Scientific | Cat#: 4385610 |
| HiSpeed Plasmid Maxi Kit | Qiagen | Cat#: 12663 |
| Lipofectamine 3000 | Thermo Scientific | Cat#: L3000008 |
| Maxima First Strand cDNA Synthesis Kit | Thermo Scientific | Cat#: K1641 |
| MEGAshortscript | Thermo Scientific | Cat#: AM1354 |
| QIAprep Spin Miniprep Kit | Qiagen | Cat#: 27106 |
| QIAquick PCR purification kit | Qiagen | Cat#: 28104 |
| Qubit RNA Broad Range Assay Kit | Thermo Scientific | Cat#: Q10210 |
| Quick-RNA Microprep | Zymo Research | Cat#: R1050 |
| Quick-RNA Miniprep | Zymo Research | Cat#: R1054 |
| SF Cell Line 4D-Nucleofector X Kit | Lonza | Cat#: V4XC-2012 |
| TGX Precast gels 12+2, 4-15% | Biorad | Cat#: 5671083 |
| TGX Precast gels 18, 4-15% | Biorad | Cat#: 5671084 |
| TGX Precast gels 26, 4-15% | Biorad | Cat#: 5671085 |
| TransBlot Turbo Midi Nitrocellulose | Biorad | Cat#: 1704159 |
| TransBlot Turbo Midi PVDF | Biorad | Cat#: 1704159 |
| TRI-reagent | Sigma-Aldrich | Cat#: T9424 |

**Supplementary Table 2:**

| **Oligonucleotides** |
|---|
| RT-qPCR primer vault RNA 1-1<br>fw: 5'-TTAGCTCAGCGGTTACTTCGACAGTTC<br>rev: 5'- AAAAGGACTGGAGAGCGCCC |
| RT-qPCR primer vault RNA 1-2<br>fw: 5'-GGCTGGCTTTAGCTCAGCGG<br>rev: 5'-AAAAGAGCTGGAAAGCACCC |
| RT-qPCR primer vault RNA 1-3<br>fw: 5'-AGCGGTTACTTCGCGTGTCATC<br>rev: 5'-AAGAGGGCTGGAGAGCGCC |
| RT-qPCR primer vault RNA 2-1<br>fw: 5'-GGGTCGGAGTTAGCTCAAGC<br>rev: 5'-AAAGGGTCAGTAAGCACCCG |
| RT-qPCR primer GAPDH<br>fw: 5'-ATGGGGAAGGTGAAGGTCG<br>rev: 5'-GGGGTCATTGATGGCAACAATA |
| RT-qPCR primer 5S<br>fw: 5'-GGCCATACCACCCTGAACGC<br>rev: 5'-CAGCACCCGGTATTCCCAGG |
| Northern blot probe for vault RNA locus 1 (not mutant sensitive):<br>5'-GAACTGTCGAAGTAACCGCTGAGCT |
| Genotyping primer vault RNA 1-1<br>fw: 5'-AAGACTCCACTCCCCTGGC<br>rev: 5'-TCCGAGGAGCCCTGATTCC |
| Sequencing primer FRT plasmid:<br>5'-TAGTTAAGCCAGTATCTGCTCC |
| Sequencing primer px458-SpCas:<br>5'-TTTATGGCGAGGCGGCGG |
| sgRNA vault RNA 1-1 #1<br>fw: 5'-CACCGCCTCAATTGTCTGGAGGTCG<br>rev: 5'-AAACCGACCTCCAGACAATTGAGGC |
| sgRNA vault RNA 1-1 #2<br>fw: 5'-CACCGCCCCGACCTCCAGACAATTG<br>rev: 5'-AAACCAATTGTCTGGAGGTCGGGGC |
| sgRNA vault RNA 1-1 #3<br>fw: 5'-CACCGAAAGGACTGGAGAGCTCCCG<br>rev: 5'-AAACCGGGAGCTCTCCAGTCCTTTC |
| sgRNA vault RNA 1-1 #4<br>fw:5'-CACCGAAAGGACTGGAGAGCGCCCG<br>rev:5'-AAACCGGGCGCTCTCCAGTCCTTTC |
